# A glial source of noradrenaline shapes synaptic integration and motor adaptation

**DOI:** 10.64898/2026.06.15.732457

**Authors:** Sambre Mach, Juliette Royer, Wenli Niu, Xia Li, Micaela Galante, Glenn Dallérac

**Affiliations:** Paris-Saclay Institute of Neurosciences, Paris-Saclay University, CNRS UMR9197, 91400, Saclay, France

## Abstract

Noradrenaline is a major neuromodulator that shapes neural circuit dynamics and behavior. Prevailing models attribute noradrenergic supply exclusively to long-range inputs from distant nuclei. Yet noradrenergic input can be weak in regions that nonetheless show marked noradrenergic modulation, suggesting additional sources. This mismatch is salient in the cerebellum, where noradrenaline influences behavior despite sparse innervation. Here, we identify Bergmann glia, radial cerebellar astrocytes, as a source of noradrenaline. Using two-photon imaging and cell-type-specific genetic manipulations, we show that Bergmann glia can synthesize and release noradrenaline through a calcium-dependent mechanism requiring the vesicular monoamine transporter VMAT2. Glia-derived noradrenaline modulates synaptic integration in Purkinje neurons and is required for motor adaptation. Together, these findings identify a glial pathway that complements the canonical long-range view of noradrenaline supply.

## Main Text

Noradrenaline (NE) is a major neuromodulator that regulates synaptic transmission, circuit dynamics, and behavior across the brain. Canonical models emphasize its release from a small set of brainstem noradrenergic nuclei, most prominently the *locus coeruleus* (LC), whose long-range axonal projections provide widespread phasic and tonic modulation of cortical and subcortical circuits, coordinating arousal, attention, and behavioral flexibility (*1–3*). Noradrenergic signaling is thus viewed as a top-down process, in which distant afferents set the neuromodulatory tone of target circuits, implicitly assuming that local neural networks play a minimal role in shaping extracellular NE levels. Yet, such an organization does not readily hold in brain regions where noradrenergic signaling exerts pronounced effects on circuit function despite sparse anatomical innervation. This mismatch is particularly salient in the cerebellum, where NE has recently been shown to markedly influence behavior (*4*, *5*), while noradrenergic fibers are sparse and heterogeneously distributed relative to the widespread expression of adrenergic receptors (*6–8*). This raises the question of whether cerebellar NE dynamics arise solely from long-range inputs or are also shaped by local mechanisms.

### Cerebellar noradrenaline arises from multiple sources

To characterize endogenous noradrenergic activity in the cerebellar cortex, we monitored spontaneous NE dynamics using two-photon imaging of the genetically encoded sensor GRAB_NE2h_ in acute cerebellar slices (*9*). Following viral delivery of the sensor to cerebellar lobules V–VII, yielding robust expression in the molecular and Purkinje cell layers (Fig. 1A, B), we detected spontaneous NE transients across all examined lobules using our event-based analysis pipeline DETECT (*10*) (Fig. 1C; Movie S1). Under baseline conditions, NE events spanned areas of 244 - 646 µm² (mean 366.5 ± 47.7 µm²), lasted 5.13 ± 0.31 s, and occurred at a frequency of 8.91 ± 2.18 min⁻¹, revealing recurrent and spatially extended noradrenergic activity (Fig. 1C-E; Movie S1, S2). Thus, although spatially localized, NE signals extended over relatively broad regions of the neuropil, consistent with extracellular diffusion. GRAB_NE2h_ selectivity was confirmed by a concentration-dependent response to NE and minimal sensitivity to dopamine (Fig. 1A, B) (*9*). Removal of extracellular Ca²⁺ nearly abolished spontaneous NE events (2.21 ± 0.48 min⁻¹), and signals recovered upon Ca²⁺ reintroduction (10.51 ± 2.13 min⁻¹; Fig. 1D, E; Movie S2), in line with an active release process (*11*).

**Fig 1.**
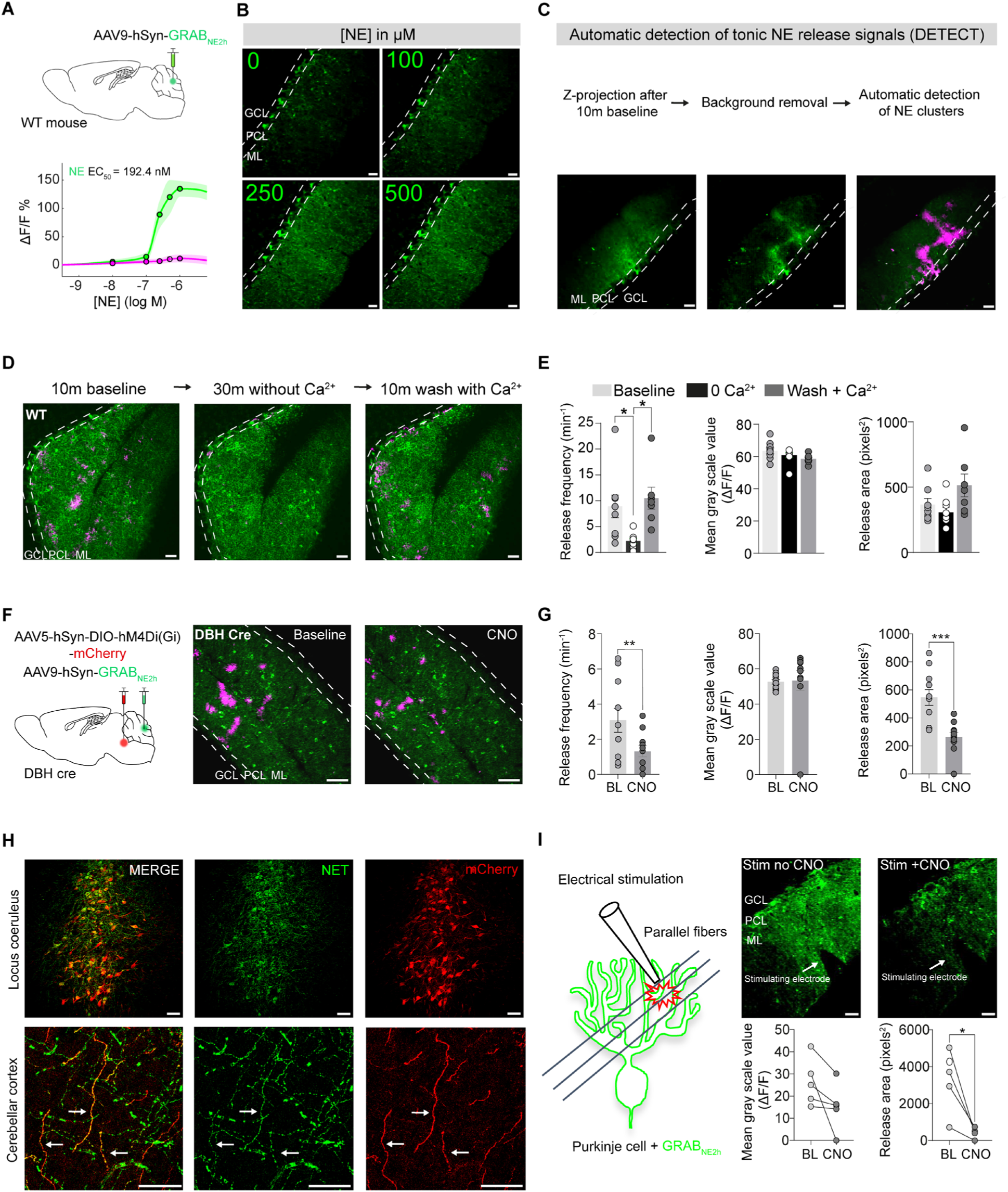
Noradrenaline release in the cerebellar cortex is not solely driven by *locus coeruleus* inputs. (**A**) Top: Schematic of AAV-GRAB_NE2h_ injection in the cerebellar cortex. Bottom: Dose-dependent GRAB_NE2h_ fluorescence increases with [NE] (n=5) or [DA] (n=4). (**B**) GRAB_NE2h_ fluorescence in the molecular layer with increasing [NE]. Scale bar, 25 µm. (**C**) Illustration of spontaneous NE release detection with the DETECT pipeline (*10*). Scale bar, 25 µm. (**D**) Temporal projections of 10 min imaging in regular ACSF, after 30 min incubation in Ca²⁺-free ACSF, and following Ca²⁺ reintroduction. Scale bar, 25 µm. (**E**) Frequency, intensity, and area of spontaneous NE events. Event frequency decreased in Ca²⁺-free conditions and recovered after Ca²⁺ reintroduction (One-way ANOVA, BL vs 0 Ca²⁺ p=0.026, Ca²⁺-free vs re-Ca²⁺ p=0.024, BL vs re-Ca²⁺ p=0.901; n=9). (**F**) Left: Schematic of viral injections in the *locus coeruleus* (LC) with the inhibitory DREADD AAV5-hSyn-DIO-hM4Di, and in the cerebellar cortex with the sensor AAV9-hSyn-GRAB_NE2h_. Middle: Temporal projections of GRAB_NE2h_ (green) recording and detected NE events (magenta). Right: Chemogenetic silencing of NE fibers with CNO (10 µM, 30 min) reduced but did not abolish NE release. Scale bar, 50 µm. (**G**) CNO significantly reduced event frequency (paired t-test, p=0.009) and area (Wilcoxon test, p=0.001; n=11). (**H**) NE fiber marker NET (green) and DREADD reporter mCherry (red) immunostaining in LC and cerebellar cortex (n=3). Scale bar, 25 µm. (**I**) Electrical stimulation evoked NE release, inhibited by chemogenetic silencing of NE fibers with CNO (paired t-test, p=0.047; n=5). GCL, granular cell layer; ML, molecular layer; PCL, Purkinje cell layer.

The prevailing model posits that NE in the cerebellar cortex originates exclusively from long-range projections arising from the LC (*12–14*). To directly assess the contribution of LC-derived inputs to cerebellar NE release, we selectively inhibited noradrenergic neurons using a chemogenetic strategy. To this end, we virally delivered a Cre-dependent inhibitory DREADD (hM4Di) to the LC of DBH-Cre mice, which express Cre recombinase in noradrenergic neurons (Fig. 1F-H), and confirmed effective silencing by the marked suppression of electrically evoked NE release in the cerebellar cortex following CNO application (Fig. 1I; Movie S3). Strikingly, chemogenetic silencing of LC noradrenergic neurons, further combined with action potential blockade by tetrodotoxin, reduced spontaneous cerebellar NE transients by only ∼50% in frequency and area (Fig. 1F,G; Movie S4). The persistence of NE signals under these conditions indicates that long-range LC projections do not fully account for cerebellar NE dynamics. Instead, a substantial fraction of spontaneous NE transients arises independently of LC input, supporting the existence of additional local sources of NE within the cerebellar cortex.

### Bergmann glia are a local source of norarenaline in the cerebellar cortex

We next sought to identify the cellular origin of this action potential-independent and LC-independent NE supply within the cerebellar cortex. Purkinje cells (PC) and Bergmann glia (BG) are both highly represented and extend elaborate processes within the molecular layer, directly overlapping areas where NE transients occur, thus positioning them as potential contributors to local NE dynamics. We therefore examined whether these cell types harbor the molecular machinery required for NE synthesis and handling. Analysis of catecholamine enzymes involved in NE synthesis showed distinct expression patterns across cell types. Tyrosine hydroxylase (TH) was densely expressed in PC across all examined lobules (Fig. 2A; Fig. S1A), consistent with a capacity for dopamine synthesis (*15*, *16*). In contrast, dopamine β-hydroxylase (DBH), the enzyme converting dopamine into NE, was strongly expressed in the majority of BG (∼75%) and largely absent from PC (Fig. 2A), pointing to BG as a cerebellar cell type capable of NE synthesis. Consistent with vesicular monoamine handling, we detected expression of the vesicular monoamine transporter VMAT2 in ∼70% of BG (Fig. 2B). Biochemical analysis of magnetically isolated astroglia confirmed VMAT2 protein expression at the expected molecular weight (60-70 kDa), and astrocyte-specific VMAT2 deletion in conditional inducible mice, hereafter termed aVMAT2cKO (*17*), reduced VMAT2 levels in BG by ∼50-60% (Fig. 2C-E). In line with a role in regulating extracellular NE availability, BG also expressed key proteins involved in monoamine transport and clearance. Although the NE transporter NET was confined to sparse LC-derived fibers and the dopamine transporter DAT, which can also transport NE, was not detected in BG (Fig. S1B, C), the high-capacity organic cation transporter OCT3, originally identified as the extraneuronal monoamine transporter (*18*, *19*), was strongly expressed in ∼75% of BG (Fig. 2B). Moreover, these glial cells also expressed the monoamine degradation enzymes MAO-B and COMT (Fig. S1D, E). Finally, to directly determine whether BG contain NE in situ, we performed NE immunostaining. NE immunoreactivity was detected within the somata and radial processes of most BG and was reduced by ∼80% in aVMAT2cKO mice (Fig. 2F,G), indicating that VMAT2 is required for NE storage in BG. NE immunoreactivity was not detected in PC (Fig. 2G), in agreement with the absence of DBH required for NE synthesis. Together, these findings indicate that BG, but not PC, are equipped with the molecular machinery required to synthesize, store, transport, and degrade NE within the cerebellar cortex.

**Fig 2.**
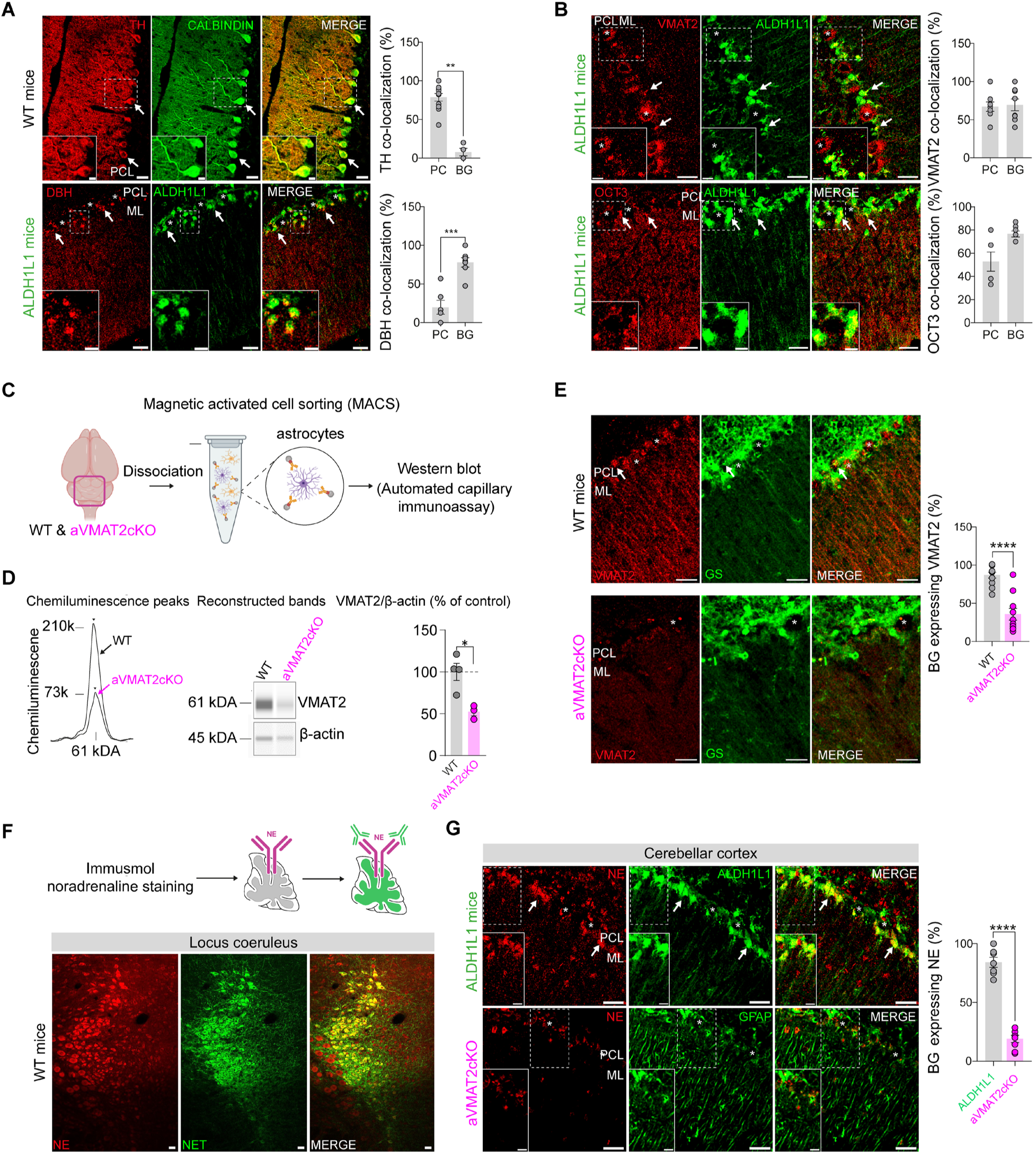
Bergmann glia harbor noradrenaline and express key monoaminergic proteins. (**A**) Immunostaining for tyrosine hydroxylase (TH, red) in Purkinje cells (PC; calbindin, green) and Bergmann glia (BG, green; ALDH1L1-eGFP mice), with quantification in PC (n=11) and BG (n=4; unpaired t-test p=0.001). Bottom: Dopamine β-hydroxylase (DBH, red) co-localization in BG (n=7) and PC (n=6; unpaired t-test p=0.0002). Scale bars, 50 µm (top), 25 µm (bottom), 10 µm (inset). (**B**) VMAT2 and OCT3 immunostaining (red) in ALDH1L1-eGFP mice (green), detected in PC and BG. Scale bar, 25 µm. (**C**) Dissociation and MACS-based isolation of cerebellar astrocytes from WT and astrocyte-specific VMAT2 conditional knockout mice (aVMAT2cKO). (**D**) *Simple Western* analysis showing reduced VMAT2 expression in aVMAT2cKO astrocytes (unpaired t-test p=0.013). (**E**) VMAT2 immunostaining (red) in WT and aVMAT2cKO mice, with astrocytes labeled by the astroglial marker glutamine synthetase (GS, green), showing reduced VMAT2-positive BG in aVMAT2cKO mice (unpaired t-test p<0.0001). Scale bar, 25 µm. (**F**) Schematic of direct NE immunostaining and control staining of *locus coeruleus* noradrenergic neurons (marker NET, green; NE, red). Scale bar, 25 µm. (**G**) Direct NE immunostaining (red) in the cerebellar cortex of control ALDH1L1-eGFP mice and aVMAT2cKO mice. BG are labeled in green by GFP or GFAP immunostaining, as indicated. NE signal in BG is reduced in aVMAT2cKO mice (unpaired t-test p<0.0001). Scale bar, 25 µm. GCL, granular cell layer; ML, molecular layer; PCL, Purkinje cell layer. Asterisks mark PC somata. Dashed lines show insets (10 µm). Arrows indicate co-localization.

To determine whether this molecular machinery translates into functional contributions to extracellular NE dynamics, we combined GRAB_NE2h_ two-photon imaging with astrocyte-specific chemogenetic manipulations. Mice were co-injected in lobules V-VII with GRAB_NE2h_ and a GFAP-driven excitatory DREADD (hM3Dq) to selectively activate BG (Fig. 3A, B). Chemogenetic activation of BG increased spontaneous NE transients’ frequency by ∼50% in wild-type mice (Fig. 3C, D; Movie S5). The efficacy of hM3Dq-mediated activation was validated in Ca²⁺ imaging experiments using GCaMP6f, which revealed an increase in astroglial Ca²⁺ transient frequency of ∼50% following CNO application (Fig. 3E, F; Movie S6). To further assess whether BG Ca²⁺ signaling is required for local NE release, we expressed the Ca²⁺ extruder CalEx (*20*) selectively in astrocytes. When combined with hM3Dq activation and GRAB_NE2h_ imaging, CalEx reduced basal NE transients’ frequency by ∼70% and abolished the hM3Dq-induced enhancement of NE release (Fig. 3D; Movie S7). In control experiments, we verified CalEx efficacy using astroglial GCaMP6f imaging during hM3Dq activation, which showed that CalEx markedly reduced the fraction of active BG and suppressed its augmentation by CNO (Fig. 3E, F; Movie S8). We next asked whether this Ca²⁺-dependent NE release requires vesicular monoamine loading in astrocytes. To this end, we used aVMAT2cKO mice, in which VMAT2 is selectively knocked down in astrocytes (Fig. 2D, E). In these mice, basal NE transient frequency in the cerebellar cortex was significantly reduced by ∼35%, and BG activation no longer enhanced NE signals (Fig. 3D), indicating that astroglial VMAT2 is required for BG-derived NE release. Finally, because PC also express VMAT2 (Fig. 2B), we also tested whether they could contribute to local NE dynamics. Selective chemogenetic activation of PC using a L7-driven excitatory hM3Dq had no effect on NE transient frequency (Fig. 3G, H), indicating that PC do not contribute to local NE release. Together, these molecular and functional findings establish BG as an active local source of NE, operating through Ca²⁺- and VMAT2-dependent mechanisms and complementing long-range LC-derived inputs.

**Fig 3.**
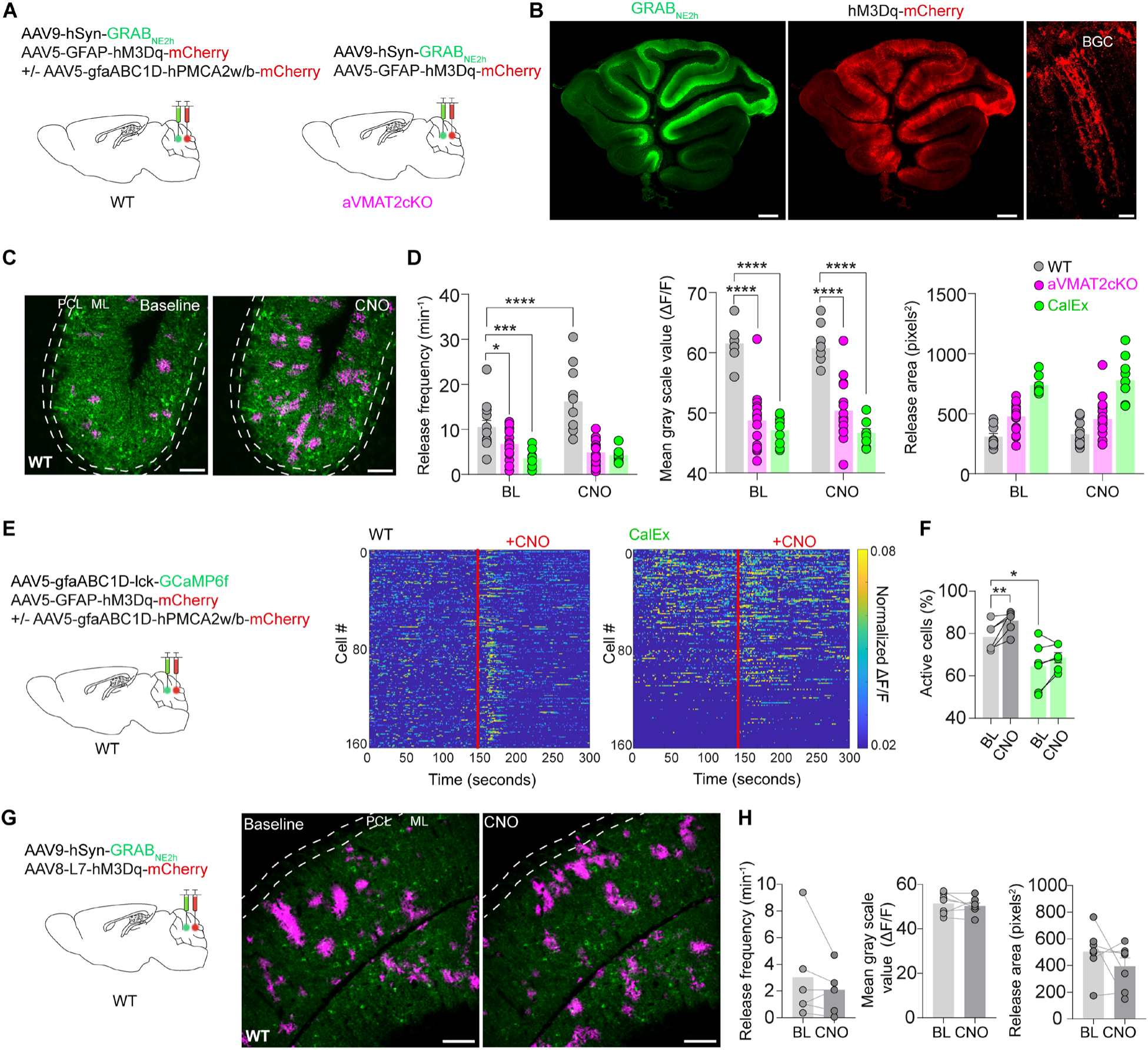
Bergmann glia drive local noradrenaline release via VMAT2. (**A**) Viral strategies to express GCaMP6f, excitatory DREADD hM3Dq-mCherry, GRAB_NE2h_, or CalEx in BG of WT and aVMAT2cKO mice. (**B**) GRAB_NE2h_ (green) and hM3Dq (red) expression in sagittal cerebellar slices (lobules IV-VII). Scale bar, 500µm. Right: zoom on BG expressing hM3Dq-mCherry. Scale bar, 25µm. (**C**) Increased spontaneous NE release after chemogenetic BG activation (CNO, 10µM) in WT mice. Scale bar, 50µm. (**D**) NE event frequency significantly increased after chemogenetic BG activation with CNO only in WT mice (two-way ANOVA, WT baseline vs CNO p<0.0001, aVMAT2cKO baseline vs CNO p=0.100, CalEx baseline vs CNO p=0.935; WT n=12, aVMAT2cKO n=19, CalEx n=6). Baseline frequency was reduced in aVMAT2cKO (two-way ANOVA, aVMAT2cKO vs WT p=0.037) and in mice harboring the calcium extruder CalEx in BG (two-way ANOVA, WT vs CalEx p<0.001). Mean event intensity was reduced at baseline in mice expressing CalEx in BG and in aVMAT2cKO mice (p<0.0001) but was unaffected by CNO. Event area was unchanged. (**E**) Astrocytic calcium imaging procedure and heatmaps showing CNO-induced chemogenetic activation in control mice but not in mice harboring CalEx in BG. (**F**) Fraction of active astrocytes increased after CNO in control mice only (two-way ANOVA; control baseline vs CNO p=0.005) and was reduced at baseline in CalEx injected mice (two-way ANOVA, control vs CalEx p=0.013; n=6 per group). (**G**) PC chemogenetic activation with the L7-driven hM3Dq did not alter NE release frequency, intensity, or area (n=7). GCL, granular cell layer; ML, molecular layer; PCL, Purkinje cell layer.

### Astroglial noradrenaline modulates synaptic inputs to Purkinje cells

NE exerts well-established modulatory effects on cerebellar circuitry, and its receptors are broadly expressed across the cerebellar cortex (*21*, *22*). Having identified BG as a local, action potential-independent source of NE, we asked how astroglial NE influences synaptic inputs onto PC. To address this, we performed *ex vivo* whole-cell patch-clamp recordings from PC while selectively activating BG using virally delivered GFAP-driven hM3Dq chemogenetic stimulation (Fig. 4A, B). Chemogenetic activation of BG increased the frequency of spontaneous excitatory postsynaptic currents (sEPSCs) by ∼30% (Fig. 4C, D) and decreased the frequency of spontaneous inhibitory postsynaptic currents (sIPSCs) by ∼35% (Fig. 4E, F). In contrast, the amplitude of neither sEPSCs nor sIPSCs was significantly altered. These effects thus produced a net shift in the balance of synaptic inputs toward excitation. Strikingly, this modulation of synaptic input frequency was completely absent in aVMAT2cKO mice, indicating that vesicular monoamine handling in BG is required for this effect (Fig. 4D, F). Likewise, pharmacological blockade of adrenergic receptors abolished the effects of BG activation on both sEPSC and sIPSC frequency, identifying NE as the functional mediator of these synaptic changes (Fig. 4D, F). Astroglial NE release therefore dynamically regulates the excitatory-inhibitory balance of synaptic inputs onto PC. Because PC constitute the sole output of the cerebellar cortex, such local astroglial control of synaptic drive most likely shapes cerebellar output and downstream motor computations (*23*). This prompted us to examine whether astroglial NE contributes to motor behavior.

**Fig 4.**
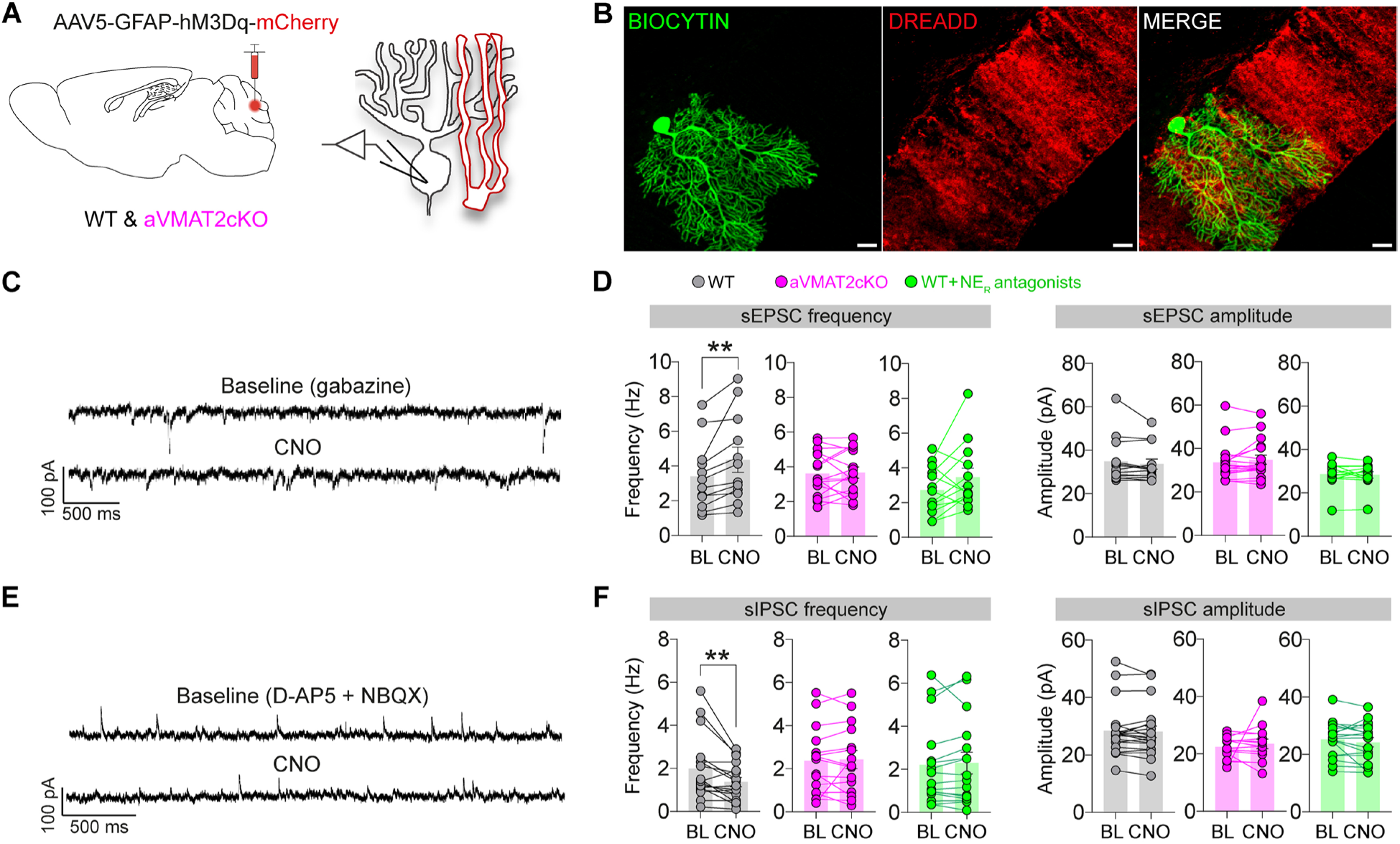
Bergmann glia-derived noradrenaline modulates Purkinje cell synaptic inputs. (**A**) Experimental design for astroglial delivery of the excitatory DREADD hM3Dq-mCherry using AAV vectors in the cerebellar cortex of WT and astrocyte-specific VMAT2 conditional knockout (aVMAT2cKO) mice. (**B**) Confocal images showing biocytin-filled Purkinje cells (PC, green) after whole-cell patch clamp and Bergmann glia (BG) expressing hM3Dq-mCherry (red) in lobule V of the cerebellar cortex. Scale bar, 20 µm. (**C**) Representative traces of spontaneous excitatory postsynaptic currents (sEPSCs) recorded from PC in WT mice under baseline conditions and following chemogenetic activation of BG with CNO (10 µM). (**D**) Quantification of sEPSC properties following BG chemogenetic activation. Mean sEPSC frequency was significantly increased by BG chemogenetic activation with CNO in WT mice (n=12; paired t-test, p=0.008) but was unchanged in aVMAT2cKO slices (n=12) or in WT slices pretreated with noradrenergic receptor antagonists (prazosin, 30 µM; propranolol, 10 µM; phentolamine, 30 µM; n=16). Mean sEPSC amplitude was unchanged across all conditions. (**E**) Representative traces of spontaneous inhibitory postsynaptic currents (sIPSCs) recorded from PC in WT mice under baseline conditions and following chemogenetic activation of BG with CNO (10 µM). (**F**) Quantification of sIPSC properties following BG chemogenetic activation. In WT mice, sIPSC frequency was significantly reduced by BG activation with CNO (n=17; Wilcoxon signed-rank test, p=0.003), an effect abolished in aVMAT2cKO mice (n=14) and in WT slices pretreated with noradrenergic receptor antagonists (n=16). Mean sIPSC amplitude was unchanged across all conditions.

### Astroglial noradrenaline regulates motor adaptation

To address the behavioral relevance of astroglial NE release in the cerebellum, we compared motor behavior in WT and aVMAT2cKO mice across tasks probing basic and adaptive locomotion while monitoring cerebellar NE dynamics *in vivo*. Basic locomotion was assessed using a treadmill paradigm that alternated self-paced locomotion with imposed belt movement requiring animals to maintain pace (Fig. S2A) (*24*). WT and aVMAT2cKO mice performed similarly across voluntary and forced locomotion (Fig. S2D, F), stopping behavior (Fig. S2E, G), and exposure to brief electrical stimuli delivered when animals failed to maintain pace (Fig. S2C), indicating preserved baseline motor performance in the absence of astroglial VMAT2. To monitor NE dynamics during this behavior, we performed miniscope imaging in mice virally transduced to express GRAB_NE2h_ in the cerebellar cortex (Fig S2B). Cranial window and baseplate implantation, followed by miniscope attachment during imaging modestly altered locomotion during treadmill on periods in both genotypes without genotype-specific effects (Fig. S2D, E). In both WT and aVMAT2cKO mice, locomotion epochs were associated with transient decreases in NE signal, whereas stop epochs were associated with gradual increases (Fig. S2H). The amplitude and duration of these locomotion- and stop-associated NE dynamics did not differ between genotypes (Fig. S2I-M), and frequency-domain analysis likewise revealed no genotype differences (Fig. S2N). Together, these data indicate that astroglial VMAT2-dependent NE signaling is not required for state-dependent NE fluctuations in the cerebellar cortex during basic locomotor behavior.

We next assessed adaptive locomotion using the horizontal ladder rung task, in which animals traversed a ladder with a regular pattern of evenly spaced rungs or an irregular pattern with missing rungs that altered stepping contingencies (Fig. 5A) (*25*). Both conditions required step-by-step control, but the irregular pattern imposed greater corrective demands and engaged adaptive recalibration. Strikingly, total stepping errors were markedly increased in aVMAT2cKO mice and remained consistently elevated across sessions (Fig. 5B, C, Movie S9). Both fine (category 1-3 on Fig. 5B) and strong (category 4-6 on Fig. 5B) errors were elevated in aVMAT2cKO mice under regular and irregular conditions (Fig. 5D, E), whereas total step number and traversal time did not differ between genotypes (Fig. S3A, B), indicating impaired adaptive control rather than reduced locomotor capacity. We then combined miniscope imaging with the ladder task (Fig. 5A) to examine NE dynamics during skilled locomotion. Cranial window and miniscope baseplate implantation did not alter step numbers or error rates in either genotype (Fig. S3A) and similarly increased traversal time in WT and aVMAT2cKO mice (Fig. S3B), while preserving the motor deficits observed in aVMAT2cKO animals (Fig. S3C). In WT mice, NE signals showed no significant change after stepping errors during regular sessions, but increased following errors during irregular sessions (Fig. 5F, G), indicating that NE responses to motor errors emerge under conditions of increased locomotor demand and variability, which require ongoing motor adaptation. In sharp contrast, error-associated increases in NE were absent in aVMAT2cKO mice, demonstrating that astroglial VMAT2-dependent signaling is required for this context-dependent response. Consistent with this, frequency-domain analysis revealed reduced NE signal fluctuations in aVMAT2cKO mice during irregular sessions, with no genotype differences under regular conditions (Fig. 5H, I).

**Fig. 5.**
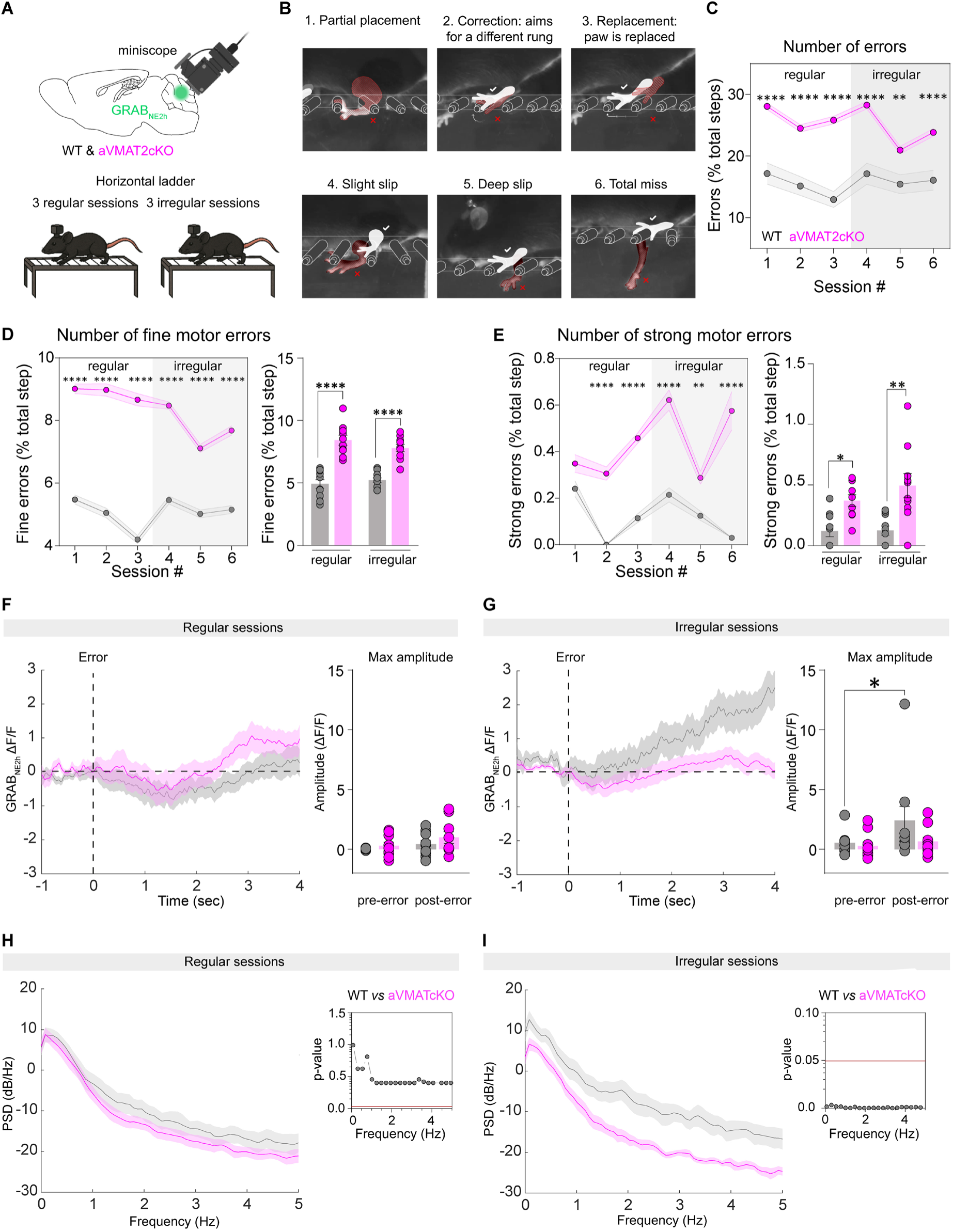
Bergmann glia noradrenaline is required for motor adaptation. (**A**) Experimental design. WT and astrocyte-specific VMAT2 conditional knockout (aVMAT2cKO) mice were injected with the GRAB_NE2h_ sensor in the cerebellar vermis and imaged with a miniscope during a horizontal ladder walking task. Animals performed sessions with regular rung spacing or irregular spacing with missing rungs. (**B**) Classification scale of fine and strong motor errors (*25*). (**C**) Total errors, expressed as percentage of steps, were increased in aVMAT2cKO mice compared to WT across sessions (two-way ANOVA, genotype effect p<0.0001). (**D**) Fine motor errors were significantly increased in aVMAT2cKO mice in both regular and irregular conditions (two-way ANOVA, genotype effect p<0.0001). (**E**) Strong motor errors were also increased in aVMAT2cKO mice compared to WT (two-way ANOVA, genotype effect p<0.0001). (**F**) Averaged GRAB_NE2h_ ΔF/F traces aligned to error onset during regular sessions showed no change in NE signal after errors in either genotype. (**G**) During irregular sessions, NE increased after errors in WT mice (two-way ANOVA, pre vs post p=0.02) but not in aVMAT2cKO mice. (**H**) Power spectral density (PSD) of GRAB_NE2h_ signals during regular sessions did not differ between genotypes. (**I**) During irregular sessions, PSD values were reduced in aVMAT2cKO mice compared to WT (frequency-wise permutation tests, p-values shown in inset). WT, n=9; aVMAT2cKO, n=10.

Together, these results indicate that astroglial NE in the cerebellum is dispensable for basic locomotion but required for adaptive motor coordination and noradrenergic dynamics evoked by motor errors during demanding tasks.

## Discussion

We here identify BG as an active source of NE that operates alongside canonical LC projections to shape cerebellar circuit function. Through a Ca²⁺- and VMAT2-dependent mechanism, astroglial NE regulates synaptic integration in PC and supports adaptive motor behavior, revealing a direct link between glial neuromodulator release, circuit computation, and behavior. These findings revise the prevailing view that noradrenergic control of brain function arises solely from long-range inputs and instead support a model in which local astroglial sources dynamically sculpt neuromodulatory tone within specific circuits. Recent studies have shown that astrocytes can act as critical mediators of noradrenergic signaling by responding to NE via their adrenergic receptors and coupling this activation to changes in synaptic and circuit function (*26–28*). Our findings go beyond this view by demonstrating that astrocytes directly provide a local source of NE within the cerebellar cortex. By coupling locally generated neuromodulator release to network demands, astrocytes can provide a mechanism for fine-scale, region-specific control alongside long-range LC-derived inputs. More broadly, identifying BG as a cellular source of a canonical monoamine suggests that glia can be integral components of neuromodulatory networks and may offer new entry points for restoring circuit function in disorders involving cerebellar dysfunction and dysregulated monoaminergic signaling.

## Acknowledgments

We thank Dr Nathalie Rouach for advice on the manuscript. We thank Julien Bouvier, Rafael Fernandes Pignoli Benzi, Andrea Giorgi and Marie Yahia for access to the horizontal ladder apparatus and guidance on its protocol.

## Funding

Fondation pour la Recherche Médicale grant FRM AJE20181039590 (GD)

Agence Nationale de la Recherche grant ANR 20-CE37-0024 (MG, GD), ANR-22-CE16-0009 (GD)

GS LSH of University Paris-Saclay as part of France 2030 programme "ANR-11-IDEX-0003 (MG)

Institut de Recherche en Santé Publique IReSP 20II134-00 (GD) and 24IReSP037IT07 (GD)

Conseil Régional d’Ile-de-France SESAME grant EX061037 (GD)

Centre Nationale de la Recherche Scientifique

Chinese Scholarship Council (XL, WN)

## Author contributions

Conceptualization: GD, MG

Methodology: SM, JR, WN, XL, MG, GD

Investigation: SM, JR, MG, GD

Visualization: SM, JR, WN, MG, GD

Funding acquisition: GD, MG, XL, WN

Project administration: GD, MG

Supervision: GD, MG

Writing - original draft: SM, GD, MG

Writing - review & editing: JR, WN, XL

## Competing interests

The authors declare no competing interests.

## Data, code, and materials availability

All data are available in the manuscript or the supplementary material. Reasonable requests for material sharing will be fulfilled upon Material Transfer Agreement.

## Supplementary Materials

### Materials and Methods

#### Animal models

Experiments were conducted on male and female mice between postnatal day 60 (P60) and P90, in accordance with the European Community Council Directives of 1 January 2013 (2010/63/EU) and with the approval of the local animal welfare committee (certificate B 91 272 108). Animals were housed in a temperature-controlled environment (25 °C) under a 12 h light/dark cycle. All mouse lines were maintained on a C57BL/6J genetic background. When no sex differences were observed, data from male and female mice were pooled for analysis. Several transgenic lines were used in this study. The ALDH1L1-EGFP line expresses the enhanced green fluorescent protein (EGFP) under the control of the astrocyte-specific Aldh1l1 promoter. The aVMAT2cKO mice were kindly provided by Paola Bezzi (University of Lausanne) and generated by crossing the GFAPcre^ERT2^ line expressing a tamoxifen (TAM) inducible cre recombinase transgene driven by the astrocytic glial fibrillary acidic protein (GFAP) promoter with the VMAT2^lox/lox^ line containing cre-excisable loxP sequences in the endogenous VMAT2 (*17*, *29*). Recombination was induced by intraperitoneal injections of tamoxifen (100 mg/kg) for five consecutive days, animals were used after a minimum of 10 days after the first injection. Finally, DBH-cre mice were kindly gifted by Bruno Giros (Paris Cité University) and express the cre recombinase transgene under the dopamine-beta-hydroxylase (DBH) promoter.

#### Immunohistochemistry

Mice were deeply anesthetized with Dolethal and transcardially perfused with iced-cold buffered saline (PBS) for exsanguination and with a 4% paraformaldehyde in PSB solution for fixation composed of (in mM): 35.4 NaH_2_PO_4_*2H_2_O, 160.7 Na_2_HPO_4_*2 H_2_O, 154 NaCl, pH 7.4. Brains were removed and post-fixed overnight at 4°C, then equilibrated with 30% sucrose at 4°C and frozen for storage at −20°C until immunohistochemistry. Sagittal 35µm thick cerebellar slices were collected using a cryostat. Brain slices were washed with PBS, permeabilized with 0.5% Triton X-100 and labeled with primary antibodies overnight at 4°C. Primary antibodies included: rabbit anti-VMAT2 (1/500, Synaptic Systems), rabbit anti-OCT3 (1/500, Gentaur), mouse anti-NET (1/500, ALOMONE AMT-002), rabbit anti-DAT (1/500, ABCAM AB184451), rabbit anti-DBH (1/500, ABCAM AB209487), mouse anti-calbindin (1/1000; Merck C9848), mouse anti-GS (1/500, Merck MABN543), rabbit anti-TH (1/500, ABCAM AB152), mouse anti-COMT (1/400, BD Biosciences #611970). Following primary antibody incubation, slices were washed twice with PBS before incubation with the appropriate secondary antibody for 2h at room temperature with the following secondary antibodies: goat anti-rabbit 555 (1/1000, Thermofischer Scientific #A-21428), goat anti-mouse 488 (1/1000, Thermofischer Scientific #A-11001), goat anti-rabbit 647 (1/1000, Thermofischer Scientific #A-21245). Slices were then washed twice with PBS and once with PB (in mM: 35.4 NaH_2_PO_4_*2H_2_O, 160.7 Na_2_HPO_4_*2 H_2_O) before being mounted in fluoromount (Thermoficher Scientific #18744094). Slices were observed under a confocal Leica SP8 microscope with ×10 (0.4 NA), x25 (0.95 NA) and x40 (1.3 NA) objectives. Images were acquired at 12 bit with identical laser power and PMT settings and analyzed with ImageJ and LasX softwares. NE staining (1/500, Immusmol #IS1028) was performed using the STAINperfect Immunostaining kit (ImmuSmol #SP-A-1000, Bordeaux, France), following the manufacturer’s protocol: mice were anesthetized and transcardially perfused with a combination of ice-cold 2% paraformaldehyde and the ImmuSmol fixation solution. Coronal and sagittal 35µm slices were collected with a cryostat (free-floating) and immunostaining was performed according to ImmuSmol protocol. Immunostainings were quantified as the proportion of cells identified by Purkinje cell or astrocytic markers that expressed the target protein.

#### Magnetic-activated cell sorting (MACS) isolation of astrocytes

Mice were deeply anesthetized with isoflurane and euthanized by cervical dislocation. The cerebellar vermis was rapidly dissected for astrocyte isolation and enzymatically dissociated at 37 °C for 30 minutes with the Miltenyi Adult Brain Dissociation Kit (130-107-677, Miltenyi Biotech, France) with gentle trituration every 10 minutes to obtain a single-cell suspension. After filtration through a through a 70 µm SmartStrainer, cells were pelleted by centrifugation (300 × g, 10 minutes, 4 °C) and resuspended in PBS + 0.5 % BSA. Fc receptors were blocked with the accompanying FcR Blocking Reagent, then cells were labeled with Anti-ACSA-2 MicroBeads (130-097-678, Miltenyi Biotech, France) for 15 minutes at 4 °C and loaded onto magnetic selection (MS) columns for positive selection. MS columns were washed twice with PBS + 0.5 % BSA and ACSA-2⁺ astrocytes eluted in 500 µL sorting buffer. For each genotype (WT and aVMAT2cKO), astrocytes isolated from five mice were pooled, counted on an automated cell counter (DeNovix, France), and pelleted by centrifugation (300 x g, 5 minutes).

#### Protein quantification by automated capillary immunoassay

Cell pellets obtained with the MACS isolation of astrocytes were lysed in 100 µL buffer containing protease and phosphatase inhibitors. Lysates were then incubated on ice for 20 minutes with intermittent mixing and cleared by centrifugation (5000 × g, 15 min, 4 °C). Protein concentrations were determined by Bradford assay (Bio-Rad). Capillary immunoassays were then performed with an “automated western blot system” (WES, ProteinSimple, BioTechne) using the 12-230 kDa Separation Module combined to either the anti-rabbit (#DM-001) or anti-mouse (#DM-00) detection modules depending on the primary antibody. This automated capillary immunodetection delivers antibody-based separation and quantification equivalent to a classical Western blot, with improved throughput and precision. Protein concentration in supernatants of isolated astrocytes was first determined by Bradford assay (Bio-Rad), adjusted to 1 µg/ml in sample buffer provided in the separation module and processed according to the protocol provided by the manufacturer. A six point calibration curve was included to determine the optimal antibody concentration for reliable and quantitative detection. Data analysis was performed using the SimpleWes software, by verifying molecular-weight markers and peak assignment. The area under the curve (AUC) for each peak was exported in excel for subsequent statistical analysis.

#### Surgery

##### Stereotaxic injections of AAVs

Mice anesthesia was induced with 4% isoflurane in 100% O_2_, and maintained with 1-1.5% isoflurane. Body temperature was monitored and maintained at 37°C during the whole procedure. The scalp was shaved and sterilized with betadine and 70% alcohol before an incision was made followed by drilling to access the skull. Intracerebral injections were achieved using a glass pipette connected to a syringe pump (Legato 130, KD Scientific). Viruses were delivered at a rate of 0.05-0.1 µL/min. Virus capillary was slowly removed after three minutes post-injection.

For the combined expression of the GRAB_NE2h_ sensor and the excitatory DREADD in WT mice, 500 nL of AAV9-hSyn-GRAB_NE2h_-GFP (WZ Biosciences Inc, 2.76×10^13^ GC/ml) and 500 nL AAV5-GFAP-hM3Dq-mCherry (Addgene, #50478, 1.8×10^13^ GC/ml) viruses were injected in lobule VI of the cerebellum (AP: −6.96 mm, ML: 0.0 mm, DV: −0.40 mm). AAV8-Pcp2-hM3Dq-mCherry viral vectors (> 4.0×10^13^ GC/ml) allowing specific excitatory hM3Dq expression in PC were provided by Viral Vector Core (University of Minnesota) and co-injected with the GRAB_NE2h_ sensor mentioned above following the same protocol. DBH-Cre mice were bilaterally injected with 200 nL of AAV9-hSyn-DIO-hM4D(Gi)-mCherry (Addgene, #44362, 2.5×10^13^ GC/ml) viral vectors in the LC (AP: −5.45 mm, ML: = +/-0.85 mm, DV = −3.65 mm) for expression of hM4Di DREADD in NE neurons. CalEx viruses (AAV5-gfaABC1D-mCherry-hPMCA2w/b, Addgene #111568, 8.9×10^12^ GC/ml) were co-injected with GFAP-hM3Dq and either GCaMP6f (AAV5-gfaABC1D-lck-GCaMP6f, Addgene #52924, 2.5×10^13^ GC/ml) or GRAB_NE2h_ in a total volume of 1 µL. All animals were used 3 to 4 weeks after injection.

##### Cranial window and baseplate implantation

For in vivo imaging, a 3 mm diameter circular craniotomy was made above the cerebellar vermis (AP: −6.5 to −7.0 mm; ML: 0.0 mm from Bregma). Three injections of GRAB_NE2h_ virus (300 nL each) in the vermis were performed before the craniotomy was sealed with a sterile glass coverslip (3 mm diameter, #1 thickness), fixed using surgical glue and by dental cement. Mice were allowed to recover for one week. Subsequently, a miniscope-compatible baseplate (Inscopix) was implanted above the cranial window using dental cement to allow miniscope fixation. Mice were then allowed to recover for an additional 2 weeks before starting imaging sessions.

#### Preparation of ex vivo cerebellar slices

Mice at P60-P80 were deeply anesthetized with isoflurane and sacrificed. The cerebellar vermis was collected after removal of the brain and placed in cold (4°C), oxygenated (95% O2 + 5% CO2) cutting artificial cerebrospinal fluid (cutting ACSF, in mM: 110 NaCl, 5 KCl, 2.5 CaCl_2_, 1.5 MgSO_4_, 1.24 KH_2_PO_4_, 10 D-glucose, 27.4 NaHCO_3_, 0.05 D-AP5). The vermis was then sagitally cut at 250 µM using a vibrating blade microtome (650HV, Microm France) for whole-cell patch clamp recordings, and at 400 µM for two-photon imaging. Slices were then transferred to room temperature recording ACSF (in mM: 110 NaCl, 5 KCl, 2.5 CaCl_2_, 1.5 MgSO_4_, 1.24 KH_2_PO_4_, 10 D-glucose, 27.4 NaHCO_3_) and allowed to recover for 1h prior to recordings.

#### Electrophysiological recordings

Individual slices were transferred to a submerged chamber mounted on a fixed-stage Scientifica SliceScope Pro 1000 electrophysiology microscope. Slices were continuously perfused with oxygenated ACSF maintained at 30-32°C with a temperature controller managed by the LinLab software (Scientifica), with a flow rate of 2mL/min. Signals were acquired with a MultiClamp 700B amplifier (Molecular Devices) filtered at 1 kHz and recordings were obtained with the Clampex software (Molecular Devices, UK). Spontaneous excitatory and inhibitory postsynaptic currents (sEPSCs and sIPSCs) were recorded from PC by whole-cell patch clamp using borosilicate glass capillaries (Harvard Apparatus,1.5 OD x 0.86 ID x 100L) with a resistance of 4-5 MΩ, filled with the following intracellular solution in mM: 144 potassium gluconate, 1 MgCl_2_, 10 HEPES, 0.5 EGTA (pH 7.4, 280 mOsm). Cells with an access resistance > 20 MΩ at resting potential were excluded from analyses as well as any cell for which a >20 % change in those parameters occurred during the course of the experiment. PC were recorded in voltage-clamp mode with a holding potential of −60 mV. Spontaneous inhibitory post-synaptic currents (sIPSCs) were recorded in the presence of the AMPA receptor antagonist NBQX (10 µM, HelloBio #HB0443) and the NMDA receptor antagonist D-AP5 (50 µM, HelloBio #HB0225) while spontaneous excitatory post-synaptic currents (sEPSCs) were recorded in the presence of the GABA_A_ receptor antagonist gabazine (5 µM, HelloBio #HB0901). Spontaneous currents were analyzed with Easy Electrophysiology software (Easy Electrophysiology Ltd, UK).

Pharmacological block of noradrenergic receptors was obtained by the application of antagonists prazosin (α1, Merck #P7791-50MG, 30 µM), propranolol (β, Merck # P0884-1G,10 µM), and phentolamine (α, Merck # P7547-100MG, 30 µM). For experiments with mice expressing DREADD, clozapine N-oxyde (CNO, Sigma-Aldrich #SML2304, 10 µM) was added to the bath solution.

In experiments using mice injected with AAV8-Pcp2-hM3Dq-mCherry, in order to confirm that patch clamped PC also expressed the excitatory DREADD, biocytin (7mg/mL, HelloBio #HB5035) was added to the intrapipette solution and slices were post-fixed overnight in 4% PFA. Slices were then washed and permeabilized before incubation with streptavidin conjugated-488 (ThermoFischer #S32354, 1/500) for 1h45. Post hoc confocal images (Leica SP8) of biocytin-filled PC and virally induced expression of mCherry were acquired to verify their colocalization.

#### Two-photon imaging

##### Acquisition

Ex vivo imaging of cerebellar slices were perfomed under a two-photon microscope (Femtonics, Budapest) equipped with an 8 kHz resonant scanner combined with a pulse laser (MaiTai-DS, SpectraPhysics, Santa Clara, CA, USA) tuned at 920 nm (green) or 1020 nm (red). Slices were continuously perfused with oxygenated ACSF (in mM: 110 NaCl, 5 KCl, 2.5 CaCl_2_, 1.5 MgSO_4_, 1.24 KH_2_PO_4_, 10 D-glucose, 27.4 NaHCO_3_) and heated at 30-32°C using an automatic temperature controller (TC-324B, Warner Instruments). Images were acquired at 31.1 Hz. Typical imaging protocol consisted in ten minutes of baseline recording followed by fifteen minutes of treatment (e.g. CNO). For evoked NE-release, a stimulating electrode (borosilicate glass capillary from Phymep (1.5 OD x 0.86 ID x 100L) 2-3 MΩ) filled with ACSF was positioned in the molecular layer of the cerebellar cortex. Stimulation was delivered using a DS2A voltage stimulator - Mk.II (Digitimer) under software control, as a single train of 20 Hz. One stimulation was acquired prior to any treatment (baseline) and CNO was bath applied for 15 min before another train of stimulation without changing the position of the stimulating electrode. Calibration-curve protocols were acquired as followed: NE or DA was applied in increasing concentrations (in µM): 0, 0.01, 0.1, 0.25, 0.5, 1, for 3 min per concentration.

##### Analysis

Videos were exported in AVI format and averaged at 9 frames per second to reduce noise and improve signal quality. Analysis was performed using a custom-built Python pipeline called DETECT (Dynamic Extraction and Tracking of Emitted Cellular Transients) (*10*), specifically designed for an automatic, GMM-based (Gaussian Mixture Model) detection of spontaneous NE or GCaMP signals depending on the experiment. This pipeline identifies individual fluorescence clusters while correcting for background noise and motion correction. Following detection, we used TrackMate in Fiji (ImageJ) to extract a range of spatiotemporal features from each identified cluster, including area, perimeter, mean intensity, total number of events, and directionality of propagation. The resulting csv files were further processed and analyzed in MATLAB using custom-written scripts for group-level statistical comparisons and visualization. For evoked NE-release, videos were exported and treated in the same way, and stimulation response area and intensity was quantified with a custom-built matlab code. Calibration-curve data were extracted as above and analyzed with a custom-built python code where fluorescence movies were processed frame-by-frame. For each frame, the mean fluorescence of non-zero pixels was calculated to exclude static background, and ΔF/F was computed using the first 60 s of the drug-free period as F₀ to minimize bleaching effects. Traces were **smoothed** with a centered moving average. Pharmacological responses were quantified by fitting ΔF/F time courses with a four-parameter logistic model using nonlinear least-squares regression. Fits typically achieved R² values of ∼0.92, and parameter stability was confirmed by sensitivity analyses with 5% added noise. Baseline values were estimated over a 180 seconds pre-stimulation period.

#### Behavioral experiments

##### Treadmill-based locomotion assessment

To assess forced locomotion, WT and aVMAT2cKO mice were tested on a 5-lane treadmill (#8710RTS Panlab Harvard Apparatus). Both groups, WT and aVMAT2cKO mice, underwent the tamoxifen injections procedure. Each treadmill lane measured 532 mm in length, 100 mm in width and 50 mm in height. The treadmill was equipped with a control unit (LE8700TS Treadmill Control Unit, Panlab Harvard Apparatus) that allowed precise adjustment of speed parameters. The protocol, adapted from (*24*), starts with a stationary period (0 cm/s) for 1 minute, followed by an initial acceleration to 10 cm/s over 30 seconds. Mice were then maintained at 10 cm/s for three intervals: from 1.5 to 2 minutes, 3 to 4 minutes, and 5 to 6 minutes, each separated by 1 minute rest phases (0 cm/s). At 7 minutes, the speed was increased to 20 cm/s for 1 minute. The total duration of the protocol was 8 minutes.

##### Horizontal ladder test

To assess skilled locomotion and sensorimotor coordination, WT and aVMAT2cKO mice were subjected to the ladder rung walking test as previously described (*25*). Both groups, WT and aVMAT2cKO mice, underwent the tamoxifen injections procedure. The apparatus consisted of two transparent plexiglass side walls (100 cm long, 20 cm high) and evenly spaced metal rungs (1 cm apart) forming a ladder elevated approximately 1 meter above the ground. Irregular sessions consisted of irregular spacing by randomly removing rungs in order to prevent pattern learning and challenge motor coordination. Each regular and irregular session consisted of 3 trials each. For irregular sessions, 3 random patterns were created and maintained identical for each mouse. Mice were familiarized to the task by performing 3 regular sessions prior to the tests. The following day, mice were tested with 3 regular sessions and 3 irregular sessions. They were placed at one end of the ladder and allowed to traverse to their home cage located at the opposite end. Crossings were recorded using a high-speed camera (200 fps, acA2040-120um, Basler Inc.) integrated into a closed-loop system that automatically adjusted the camera position to precisely track the mice as they traversed the ladder. The camera was positioned laterally and slightly inclined to capture precise paw movements. Videos were analyzed frame-by-frame (MPC software) to evaluate forelimb and hindlimb placements using a 7-points foot fault scoring system (*25*):

- 0: correct placement
- 1: partial placement, paw is misplaced with only toes or heels
- 2: correction, paw aims for one rung but is placed on another without touching the first
- 3: replacement, paw position is adjusted after first initial placement
- 4: slight slip without fall
- 5: deep slip, paw slips leading to imbalance and fall
- 6: total miss, paw misses the rung leading to a fall.

For analysis, errors were expressed as a percentage of total steps to account for potential differences in step count across trials. The total number of errors per session and the corresponding error score intensity were measured.

#### GRAB_NE2h_ miniscope imaging

To control for potential bias from the surgery or miniscope attachment on motor performances, mice performed the treadmill and horizontal ladder protocol before and after surgery for GRAB_NE2h_ imaging. Imaging acquisition during the task was performed using the nVista miniscope system (Inscopix), fixed on the surgically implanted baseplate, with a 20 frames per second acquisition rate. Raw signals were then processed in Inscopix Data Processing Software as followed: after motion correction, regions of interest (ROIs) were manually defined based on signal quality while avoiding blood vessels, using the same ROI size across videos. ΔF/F traces were then extracted and normalized for each ROI by their root mean square, and aligned to the behavioral videos for correlation analyses. Correlation analyses, power spectral density estimates, and statistical tests were performed in MATLAB.

#### Statistics

All data were analyzed using GraphPad Prism, MATLAB, and Python. Data normality was assessed using the Shapiro-Wilk test. Outliers were identified using the Robust Regression and Outlier Removal (ROUT) method (Q = 1%) (*30*) implemented in GraphPad Prism, which controls the false discovery rate, and were removed prior to statistical analysis. For comparisons between two groups, two-tailed paired or unpaired Student’s t tests were used when data were normally distributed. Otherwise, the Wilcoxon signed-rank test (paired) or Mann-Whitney U test (unpaired) was applied. One- or two-way ANOVA, including repeated-measures designs where appropriate, were used, followed by post hoc multiple-comparison tests using Tukey’s or Sidak’s methods as appropriate. All data are reported as mean ± SEM. Statistical significance was set at α = 0.05. *p < 0.05, **p < 0.01, ***p < 0.001 and ****p < 0.0001.

**Fig. S1.**
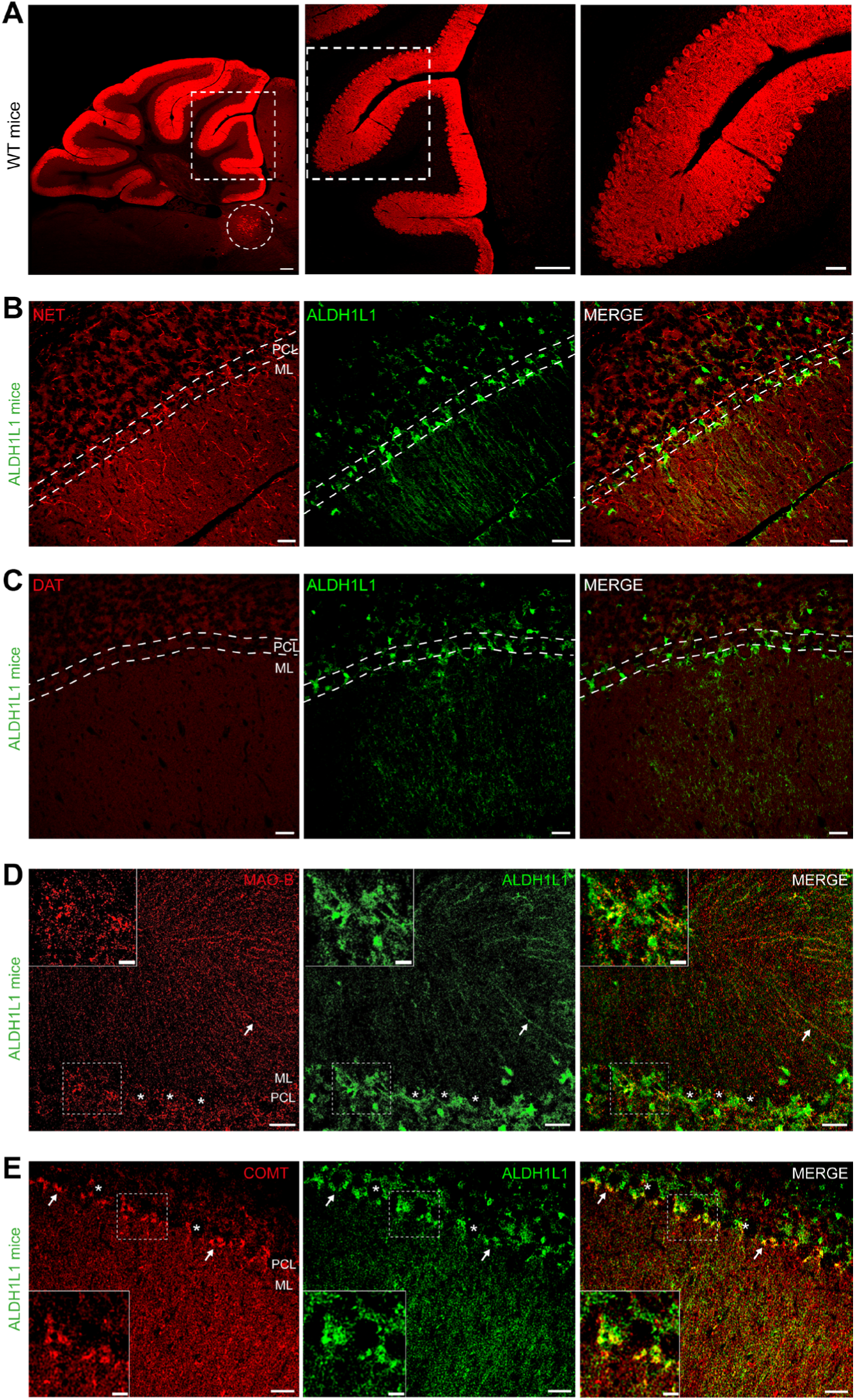
Monoaminergic signaling components in the cerebellar cortex. (**A**) Tyrosine hydroxylase (TH, red) immunostaining in WT mice showing widespread labeling across cerebellar lobules. Dashed squares indicate regions shown at higher magnification (middle and right panels). The dashed circle highlights TH-positive *locus coeruleus* neurons (n = 8). Scale bar, 100 µm. (**B**) NET immunostaining (red) in ALDH1L1-eGFP mice (green) showing NET expression restricted to sparse fibers within the molecular layer and absent from Bergmann glia (BG). Scale bar, 25 µm (n=4). Dashed lines delineate the Purkinje cell layer (PCL) and molecular layer (ML). (**C**) DAT immunostaining (red) in ALDH1L1-eGFP mice (green) showing absence of DAT expression in the cerebellar cortex. Scale bar, 25 µm (n=2). Dashed lines delineate the Purkinje cell layer (PCL) and molecular layer (ML). (**D**) MAO-B immunostaining (red) in ALDH1L1-eGFP mice (green) showing expression of monoamine degradation enzymes in BG. Insets show higher magnification views. Scale bars, 25 µm (main), 10 µm (inset) (n=2). (**E**) COMT immunostaining (red) in ALDH1L1-eGFP mice (green) confirming expression of monoamine degradation pathways in BG. Insets show higher magnification views. Scale bars, 25 µm (main), 10 µm (inset) (n=3).

**Fig. S2.**
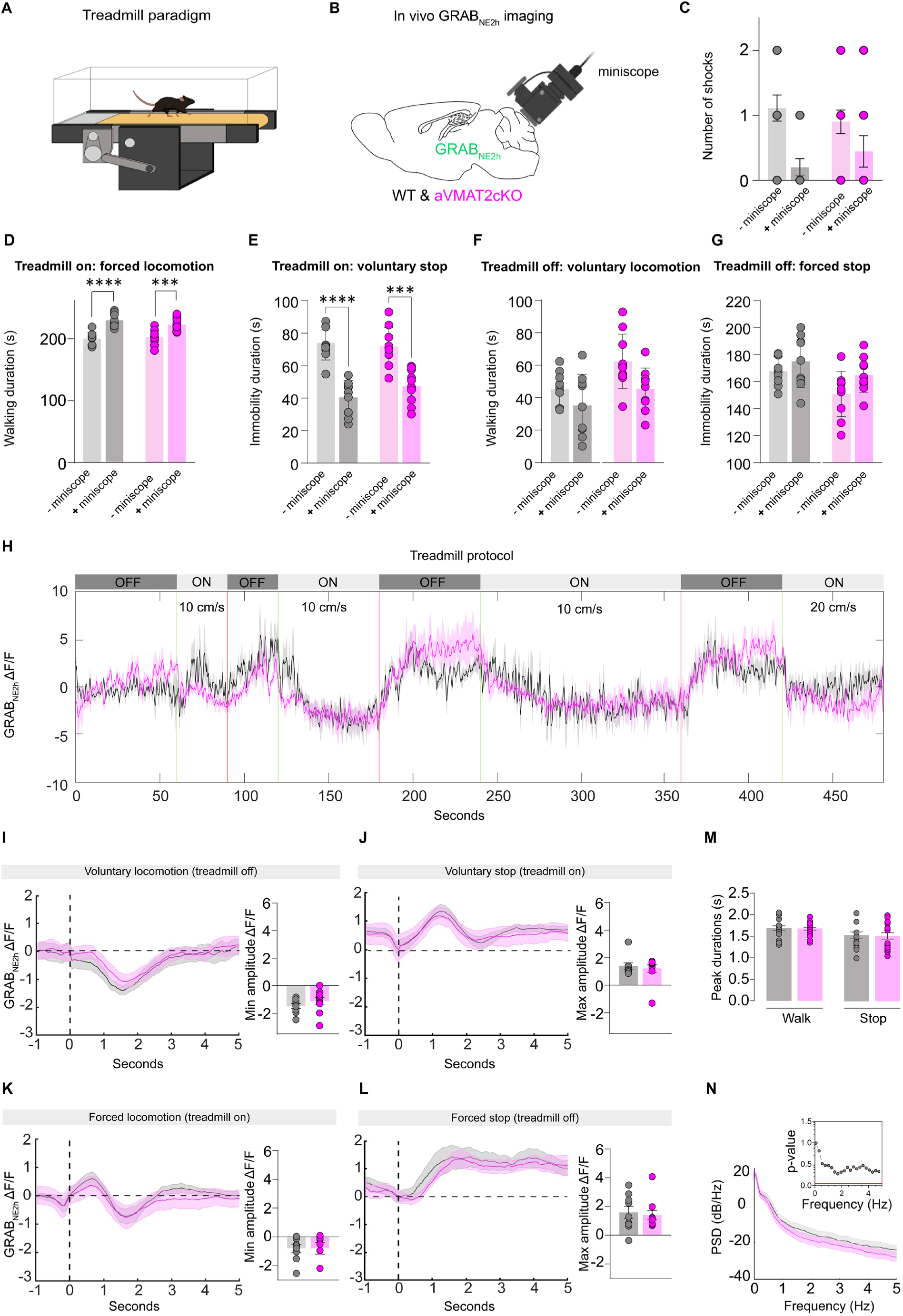
Basal locomotion and associated NE dynamics are preserved after VMAT2 deletion in BG. (**A**) Schematic of the treadmill apparatus used for locomotion experiments. The light gray area corresponds to the region where mice receive a shock if they fail to keep pace. (**B**) Schematic of miniscope placement for GRAB_NE2h_ imaging in the cerebellar cortex. (**C**) The number of electrical shocks delivered when mice fail to keep pace during treadmill ON periods is low and not significantly different with or without miniscope (2-way ANOVA, p=0.48) or between WT and aVMAT2cKO mice (2-way ANOVA, p=0.37). (**D**) Walking duration during treadmill ON periods (“forced locomotion”) is increased when the miniscope is attached (2-way ANOVA, p<0.001 for both WT and aVMAT2cKO) but does not differ between WT and aVMAT2cKO mice (2-way ANOVA, p=0.49). (**E**) Immobility duration during treadmill ON periods (“voluntary stop”) is reduced when the miniscope is attached (2-way ANOVA, p<0.001 for both WT and aVMAT2cKO) and does not differ between WT and aVMAT2cKO mice (2-way ANOVA, p=0.49). (**F**) Walking duration during treadmill OFF periods (“voluntary locomotion”) is unchanged when the miniscope is attached (2-way ANOVA, p=0.50 for WT and p=0.08 for aVMAT2cKO) and does not differ between WT and aVMAT2cKO mice (2-way ANOVA, p=0.09 without miniscope, p=0.47 with miniscope). (**G**) Immobility duration during treadmill OFF periods (“forced stop”) is unchanged when the miniscope is attached (2-way ANOVA, p=0.76 for WT and p=0.19 for aVMAT2cKO) and does not differ between WT and aVMAT2cKO mice (2-way ANOVA, p=0.09 without miniscope, p=0.47 with miniscope). (**H**) Treadmill protocol with corresponding mean GRAB_NE2h_ ΔF/F traces for WT (black) and aVMAT2cKO (magenta) mice. Green and red vertical lines indicate treadmill OFF-to-ON and ON-to-OFF transitions, respectively. (**I**) Mean GRAB_NE2h_ ΔF/F signals aligned to voluntary locomotion onset (treadmill OFF) show similar ΔF/F decrease between WT and aVMAT2cKO mice (Mann-Whitney U test, p=0.16). (**J**) Mean GRAB_NE2h_ signals aligned to voluntary stop onset (treadmill ON) show similar ΔF/F increase between genotypes (Mann-Whitney U test, p=0.18). (**K**) Mean GRAB_NE2h_ signals aligned to forced locomotion onset (treadmill ON) show no genotype differences in ΔF/F decrease (Mann-Whitney U test, p=0.50). (**L**) Mean GRAB_NE2h_ signals aligned to forced stop onset (treadmill OFF) show similar ΔF/F increase between genotypes (Mann-Whitney U test, p=0.78). (**M)** Peak duration of NE transients during locomotion and stop epochs is comparable between WT and aVMAT2cKO mice (locomotion: unpaired t-test, p=0.82; immobility: unpaired t-test p=0.88). (**N**) Power spectral density (PSD) analysis of NE signals shows no differences across frequencies between genotypes (frequency-wise permutation tests, p-values shown in inset). WT, n=9; aVMAT2cKO, n=10.

**Fig. S3.**
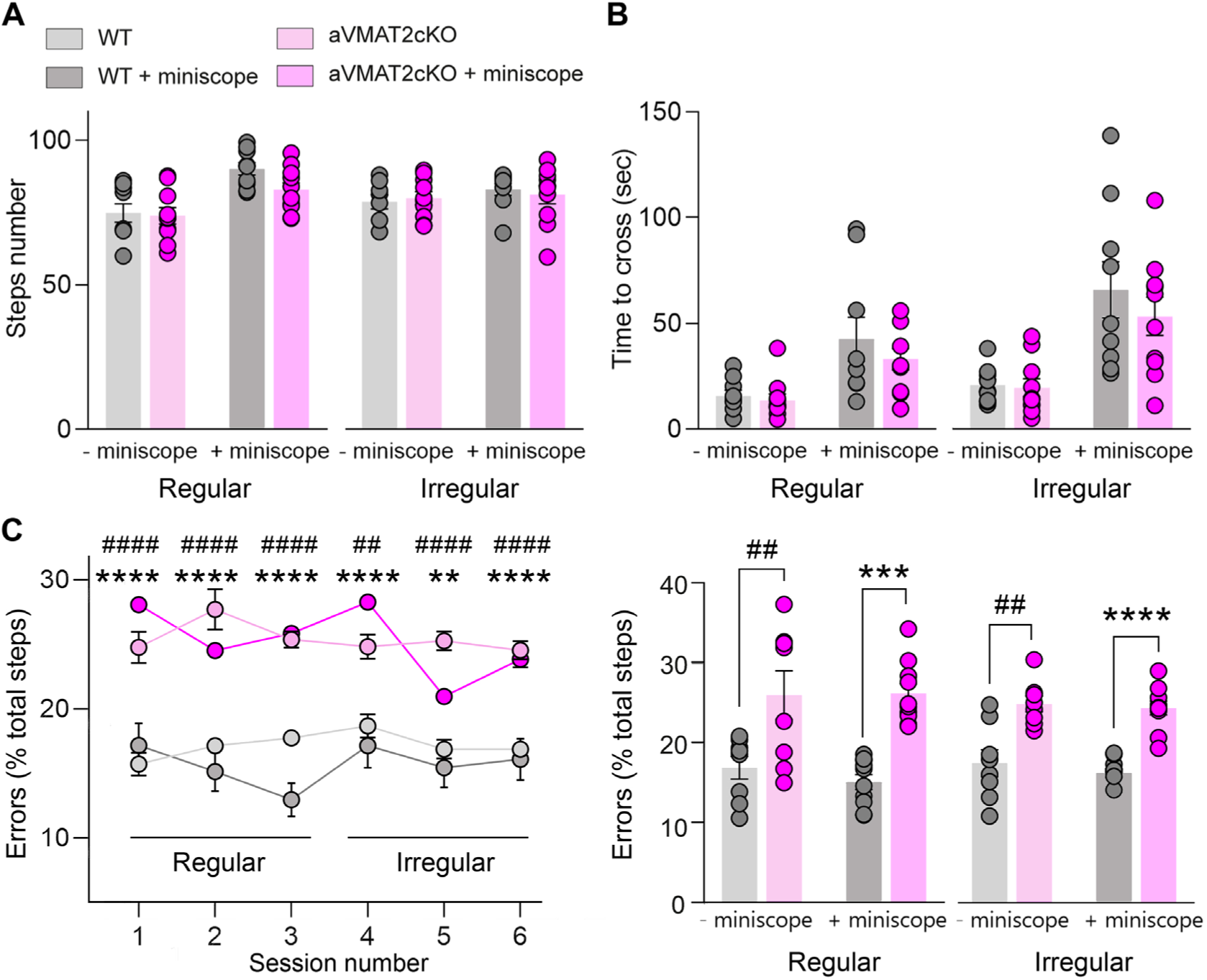
Horizontal ladder motor phenotypes are preserved after miniscope attachment. (**A**) Total number of steps during ladder traversal in WT and aVMAT2cKO mice under regular and irregular conditions shows no genotype differences without (2-way ANOVA, regular p=0.99, irregular p=0.99) and with (2-way ANOVA, regular p=0.69, irregular p=0.71) miniscope. (**B**) Time to cross the ladder is comparable between WT and aVMAT2cKO mice under both regular and irregular conditions, without (2-way ANOVA, regular p=0.99, irregular p=0.98) and with (2-way ANOVA, regular p=0.24, irregular p=0.96) miniscope. (**C**) Left: Stepping error rate (% of total steps) across sessions under regular and irregular conditions. Right: Stepping error rates (% of total steps) are unchanged by miniscope attachment in both WT (2-way ANOVA, regular p=0.41, irregular p=0.12) and aVMAT2cKO mice (2-way ANOVA, regular p=0.685, irregular p=0.716). The increase in error rate in aVMAT2cKO remains significant without (2-way ANOVA, regular p=0.007, irregular p=0.0002), or with (2-way ANOVA, regular p=0.0002, irregular p<0.0001) miniscope. WT, n=9; aVMAT2cKO, n=10. Hash symbols (#) indicate genotype comparisons between WT and aVMAT2cKO mice without miniscope attachment, whereas asterisks (*) indicate genotype comparisons between WT and aVMAT2cKO mice with miniscope attachment. WT, n = 9; aVMAT2cKO, n = 10.

## Movie captions

**Movie S1. Detection of spontaneous NE transients in the cerebellar cortex.**

Left: Raw two-photon GRAB_NE2h_ imaging showing spontaneous NE transients. Middle: Background removal using our custom pipeline (DETECT) (*10*), isolating spontaneous NE transients. Right: Detection and segmentation of NE transients (magenta) for quantitative analysis. Scale bar, 25 µm. Displayed at 20x real-time speed.

https://doi.org/10.5281/zenodo.20706364

**Movie S2. Extracellular Ca²⁺ removal abolishes spontaneous NE release.** Left: Two-photon GRAB_NE2h_ imaging of spontaneous NE transients under baseline conditions in the cerebellar cortex. Right: Two-photon imaging of spontaneous NE transients after extracellular Ca²⁺ removal. Scale bar, 25 µm. Displayed at 20x real-time speed.

https://doi.org/10.5281/zenodo.20706439

**Movie S3. Chemogenetic silencing of LC noradrenergic neurons suppresses evoked NE transients in the cerebellar cortex.** Left: Two-photons GRAB_NE2h_ imaging of electrically evoked NE release in the molecular layer under baseline conditions. Right: Electrical stimulation after hM4Di chemogenetic silencing of LC noradrenergic neurons with CNO (10 µM, 30 min), showing suppression of evoked NE transients. Scale bar, 25 µm. Displayed at 20x real-time speed.

https://doi.org/10.5281/zenodo.20706474

**Movie S4. Chemogenetic silencing of LC noradrenergic neurons does not abolish spontaneous NE release.** Left: Two-photon GRAB_NE2h_ imaging of spontaneous NE release under baseline ACSF conditions. Right: Two-photon GRAB_NE2h_ imaging following chemogenetic silencing of LC noradrenergic neurons with CNO (10 µM, 30 min), showing persistent spontaneous NE release. Scale bar, 25 µm. Displayed at 20x real-time speed.

https://doi.org/10.5281/zenodo.20706542

**Movie S5. Chemogenetic activation of Bergmann glia increases spontaneous NE release.** Left: Two-photon GRAB_NE2h_ imaging of spontaneous NE transients under baseline ACSF conditions in the cerebellar cortex. Right: Two-photon GRAB_NE2h_ imaging during hM3Dq chemogenetic activation of Bergmann glia with CNO (10 µM), showing increased spontaneous NE transients. Scale bar, 25µm. Displayed at 20x real-time speed.

https://doi.org/10.5281/zenodo.20706602

**Movie S6. Chemogenetic activation increases Bergmann glia Ca²⁺ dynamics.** Left: Two-photon GCaMP6f imaging of Bergmann glia Ca²⁺ transients under baseline ACSF conditions. Right: Two-photon GCaMP6f imaging during astroglial hM3Dq chemogenetic activation with CNO (10 µM), showing increased astroglial Ca²⁺ dynamics. Scale bar, 25 µm. Displayed at 5x real-time speed.

https://doi.org/10.5281/zenodo.20706651

**Movie S7. Disruption of Bergmann glia Ca²⁺ signaling reduces spontaneous NE transients and suppresses hM3Dq-induced increases in spontaneous NE release.** Left: Two-photon GRAB_NE2h_ imaging of spontaneous NE release under baseline ACSF conditions in mice expressing the calcium extruder CalEx in astroglial cells. Right: Two-photon GRAB_NE2h_ imaging during chemogenetic activation of Bergmann glia with CNO (10 µM) in mice expressing CalEx in astroglial cells, showing suppression of the hM3Dq-induced increase in spontaneous NE release. Scale bar, 25 µm. Displayed at 20x real-time speed.

https://doi.org/10.5281/zenodo.20706691

**Movie S8. CalEx suppresses Bergmann glia Ca²⁺ dynamics during chemogenetic activation.** Left: Two-photon GCaMP6f imaging of Bergmann glia Ca²⁺ transients under baseline ACSF conditions in mice expressing the calcium extruder CalEx in cerebellar cortex astrocytes. Right: Two-photon GCaMP6f imaging showing reduced Bergmann glia Ca²⁺ dynamics during astroglial chemogenetic activation with CNO (10 µM) in mice expressing CalEx in astrocytes. Scale bar, 25 µm. Displayed at 5x real-time speed.

https://doi.org/10.5281/zenodo.20706743

**Movie S9. aVMAT2cKO mice show impaired motor coordination during the horizontal ladder task.** Top: WT mouse performing the horizontal ladder task under irregular conditions. Bottom: aVMAT2cKO mouse performing the same task, showing increased fine and strong stepping errors. Displayed at 1x real-time speed.

https://doi.org/10.5281/zenodo.20706802

## Notes

### Competing Interest Statement

The authors have declared no competing interest.

